# Non-Invasive Brain Stimulation Data Analysis Structure (NIBS-DAS): A Template for the Layout, Management, and Analysis of NIBS Data

**DOI:** 10.64898/2026.04.30.720417

**Authors:** Michael P. Barham, Jordan Morrison-Ham, Christopher J. Greenwood, Giacomo Bertazzoli, Nigel C. Rogasch, Hannah G. K. Bereznicki, Ellen F. P. Younger, Elizabeth G. Ellis, Liam G. Graeme, David A. Cunningham, Wei-Yeh Liao, Peter J. Fried, Alvaro Pascual-Leone, Peter G. Enticott, Daniel T. Corp

## Abstract

Currently, there is no consensus about how investigators should format their NIBS data for sharing. This presents a barrier to the advancement of big data analyses because it requires time-consuming operations to generate consistent formats across different shared datasets. Recently, we launched ‘Big non-invasive brain stimulation data’ (Big NIBS data), an open-access platform and repository for NIBS data (*https://www.bignibsdata.com/),* providing a structured mechanism for researchers to share NIBS data. However, the reusability and interoperability of data uploaded to Big NIBS data is restricted by the absence of a common data structure. The current paper addresses this problem by creating the ‘NIBS data analysis structure’ (NIBS-DAS), a template pipeline for the layout, management, and analysis of collated NIBS outcome data. While its primary purpose is to provide a template layout for uploading collated data to the Big NIBS data repository, NIBS-DAS also offers guidelines for the management and analysis of collated NIBS data, thereby forming a data analysis pipeline that can be freely used by the NIBS field in general. We anticipate that NIBS-DAS will serve to facilitate data sharing on the Big NIBS data platform and promote greater standardisation of data management and analytical practices in the NIBS field.

## Introduction

Non-invasive brain stimulation (NIBS) refers to a suite of techniques used clinically to modulate brain networks in psychiatric and neurological disorders, and experimentally to probe the activity of the nervous system (Bhattacharya et al., 2022; Polanía et al., 2018). One of the major issues in clinical and neurophysiological NIBS studies is the variability of intraindividual and interindividual response (Corp et al., 2020, 2021; López-Alonso et al., 2014, 2015). Small sample sizes inherent to many single-site studies have hampered our ability to demonstrate reproducible NIBS effects, and sources of response variability (Corp et al., 2020, 2021; Guerra et al., 2020a, 2020b). To this end, we recently launched ‘Big NIBS data’ (Corp et al., 2025), the first NIBS open-access platform and repository, to increase the FAIRness (Findability, Accessibility, Interoperability, Reusability) of NIBS data and promote big data analyses in the field (Wilkinson et al., 2016), and in turn provide better insights into the clinical efficacy of NIBS protocols, sources of intra and inter-individual variability, and electrophysiological abnormalities in brain disorders, among other possibilities.

Big NIBS data (Corp et al., 2025) allows the upload of both raw and collated NIBS data. In the present context, ‘raw data’ refers to data in their original form prior to processing (e.g., machine-generated outputs like raw EMG traces), ‘uncollated data’ refers to data, whether raw or processed, that are not yet organised in a format suitable for statistical analysis, while ‘collated data’ refers to processed data in which variables (dependent variables [DVs], independent variables [IVs], and covariates) and observations are organised in a tabular structure suitable for downstream data management and analysis. The upload of raw data to Big NIBS data is important for re-use (e.g., to extract alternative outcomes later) and transparency. Yet, collated NIBS data is a critical focus of Big NIBS data because it greatly expedites data sharing and re-use; sharing raw data only would mean that each user would need to extract and collate the data independently, resulting in duplication of effort.

However, at present, the reusability and interoperability of data uploaded to Big NIBS data is restricted by the absence of a common collated data structure. Therefore, integration of NIBS datasets currently requires manual and impromptu reformatting prior to analyses. Greater uniformity in dataset layout would streamline this process on the Big NIBS data repository and move pooled analyses of NIBS datasets beyond small-scale collaborations. In addition, there is a paucity of standardised guidelines for data handling in the NIBS field. Practices such as how NIBS data are managed to ensure reproducibility of results, and guidance for statistical analyses, are not yet formalised in the field. Greater standardisation would reduce variability produced by differences in data handling procedures between NIBS laboratories.

One of the most widely used systems for data management in neuroscience is the Brain Imaging Data Structure (BIDS) (Gorgolewski et al., 2016). BIDS has now been extended from MRI to MEG (Niso et al., 2018), EEG (Pernet et al., 2019), genetics (Moreau et al., 2020), PET (Norgaard et al., 2022) and microscopy (Bourget et al., 2022) among others. Investigators are also currently working on a BIDS extension proposal for NIBS (NIBS-BIDS; Bertazzoli et al., 2026). Consistent with the broader BIDS framework, NIBS-BIDS focuses on standardising the structure and description of raw, machine-generated NIBS data. However, BIDS and NIBS-BIDS were not designed to address later stages of the data lifecycle, such as how raw data should be collated into statistical analysis-ready tabular formats, managed for reproducibility, prepared for sharing with collaborators (e.g., biostatisticians), or integrated with other NIBS datasets for big data analyses. Guidelines for these downstream steps are therefore still needed.

The aim of this paper is to meet this need by creating the ‘non-invasive brain stimulation data analysis structure’ (NIBS-DAS): a template for the layout, management, and analysis of collated NIBS outcome data. The primary purpose of NIBS-DAS is to provide a common, standardised layout for uploading collated data to the Big NIBS data repository, enabling greater reusability and interoperability. In addition, given the absence of such advice in the field, NIBS-DAS also provides guidelines for the management and analysis of this collated data, forming a template pipeline for data analysis that can be freely adopted by the NIBS field in general.

Importantly, NIBS-DAS was designed to be complementary to NIBS-BIDS, allowing future integration into a more complete data acquisition-to-analysis pipeline. The definition of these structures becomes particularly important with the emerging potential for automated NIBS data pipelines. In such a framework, raw datasets could be structured and described in NIBS-BIDS format and then collated by automated software into NIBS-DAS format for statistical analysis and further sharing. In this context, NIBS-DAS provides a target collated data schema for automated pipelines. Researchers from NIBS-BIDS have actively collaborated on the present project to ensure alignment of these aims.

Given there are numerous ways that one could appropriately handle NIBS data across different types of NIBS studies and objectives, we do not propose NIBS-DAS as the only way to layout, manage, and analyse collated NIBS data. Instead, we create NIBS-DAS as a logical template that can be adapted to different contexts.

Here, we provide two example datasets, with code files for data management and statistical analysis, to demonstrate how NIBS-DAS supports common analytical workflows. The present paper constitutes version 1.0 of NIBS-DAS. All data and code described below can be found at *https://github.com/danieltcorp/nibs-das_repo* (Corp & Barham, 2026), along with future updates.

## Methods and Results

NIBS-DAS is divided into three steps: data layout; data management; and data analysis (Figure 1). Each step of our pipeline is described below, and two simulated datasets from common types of NIBS experiments are provided on which we demonstrate the NIBS-DAS pipeline. The first dataset is an entirely simulated clinical trial using repetitive transcranial magnetic stimulation (rTMS) as a treatment for major depressive disorder, with symptom scores as the DV. The second dataset is a simulated neurophysiological experiment assessing corticospinal excitability following single and paired-pulse TMS, with motor-evoked potential (MEP) amplitudes as the DV. Given there are numerous other NIBS techniques, such as transcranial electrical stimulation (tES) and transcranial ultrasound (TUS), these publicly available simulated datasets and the accompanying data management and analysis code are intended to serve as templates which researchers can adapt to suit their projects.

**Figure 1.**
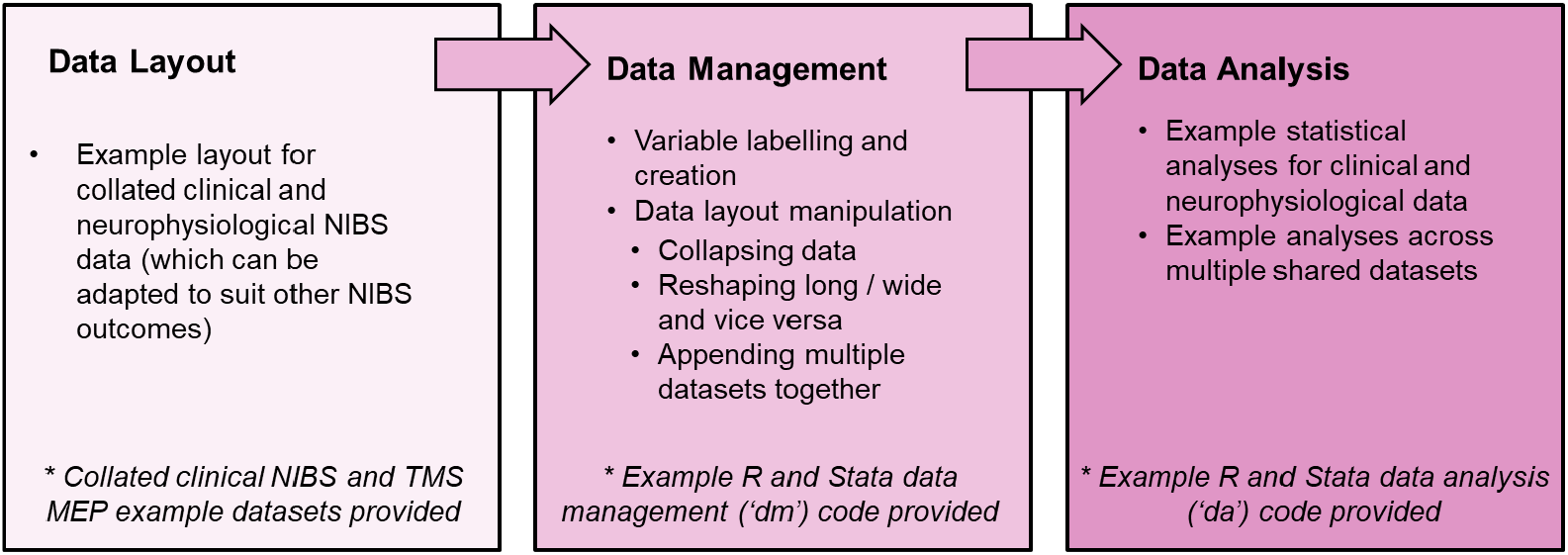
The three components of NIBS-DAS: data layout, data management, and data analysis.

These datasets are organised within a template NIBS-DAS folder structure for both simulated experiments. This folder structure is shown in Figure 2 to provide context to the operations performed on these files, and is designed to facilitate file organisation, tracking of operations, and version control of NIBS experiments. Further elaboration of the folder structure and the handling of the files within them are provided in the README document on our GitHub repository.

**Figure 2.**
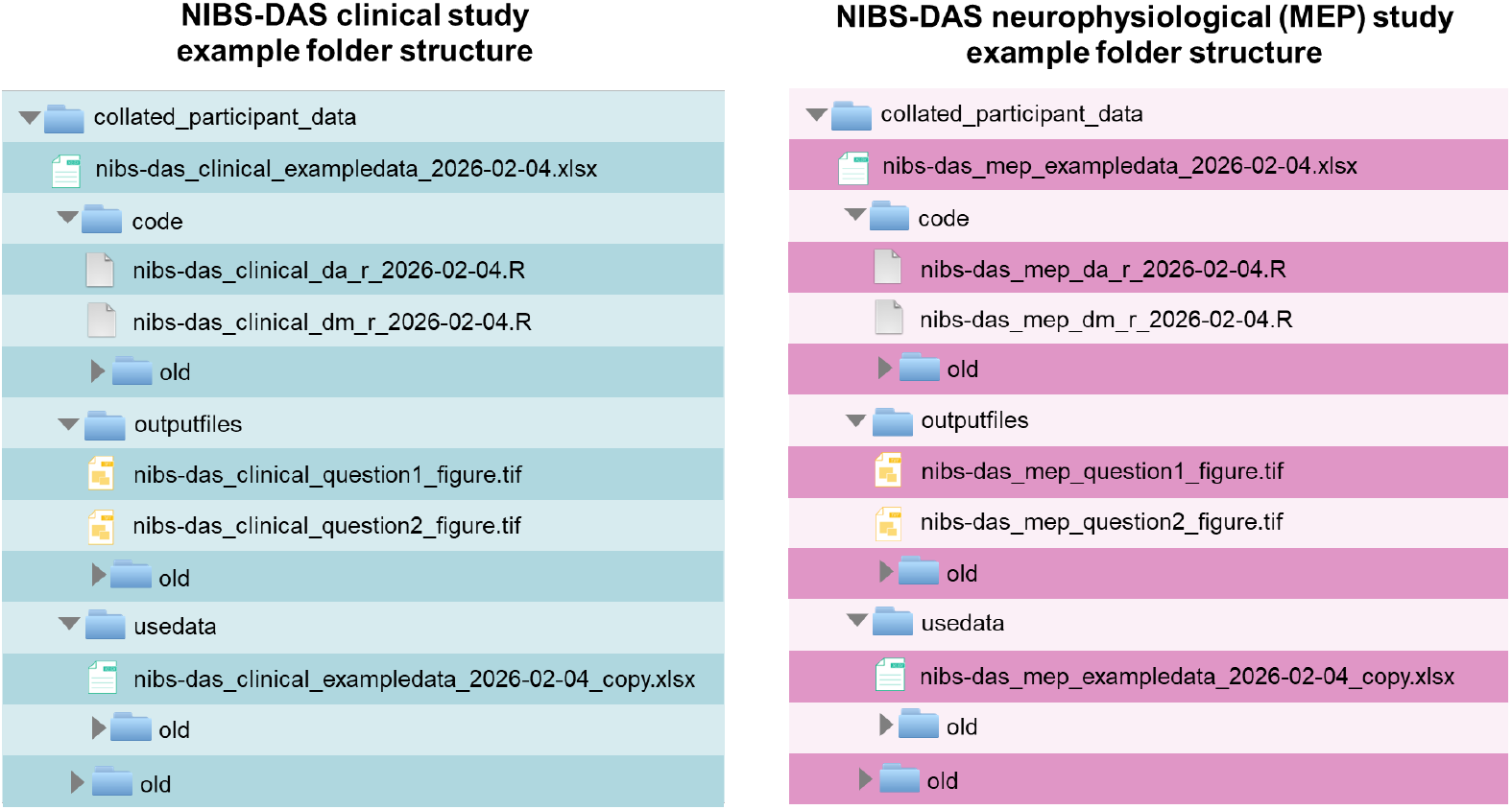
Example folder structure used to organise and manage collated data from our example clinical and neurophysiological (measuring MEPs) experiments. NIBS-DAS folders and files are named using snake_case and dated using the YYYY-MM-DD format to facilitate version control.

While not the explicit aim of the present paper, we also provide the corresponding folders containing the raw data for these example clinical and neurophysiological (MEP) experiments at: *https://github.com/danieltcorp/nibs-das_repo/tree/main/raw_data_examples*, based primarily on current NIBS-BIDS (Bertazzoli et al., 2026) recommendations, with additional guidance from EMG-BIDS (*https://bids-specification.readthedocs.io/*). This provides users with context for how NIBS-DAS fits within a complete dataset, how data can be collated into NIBS-DAS format from the original raw data (in NIBS-BIDS format), and how these two formats can function together. Importantly, this also provides template metadata files, which should be submitted alongside any data uploaded to the repository at *https://www.bignibsdata.com/* to describe the content of the dataset/s and the methods used to collect them. The aim of NIBS-DAS is not to provide guidance on metadata formatting, as this would duplicate the efforts of NIBS-BIDS. Our metadata files are formatted using the current version of NIBS-BIDS (v6); yet are only templates based on our simulated experiments. We recommend that users follow NIBS-BIDS (Bertazzoli et al., 2026) and FAIR principles (Wilkinson et al., 2016), when creating metadata files, and tailor these to data contained within their own NIBS experiments.

### Data Layout

The NIBS-DAS pipeline starts from a spreadsheet at the root directory (Figure 2), containing participant and study data collated from raw data. We follow BIDS conventions for variable naming. In BIDS, participant-level variables in TSV files are typically formatted in snake_case, whereas study-level metadata fields are defined in accompanying JSON files (e.g., ‘ManufacturerModelName’) using camelCase/PascalCase. In NIBS-DAS, both participant and study-level data are collated into a single spreadsheet to enable a wider range of big-data analyses (e.g., the effect of different machines across pooled datasets). Therefore, in NIBS-DAS, these study-level variable names are preserved and imported to the spreadsheet exactly as they appear in the original metadata files (e.g., ‘ManufacturerModelName’).

How data are organised within a spreadsheet influence how it can be subsequently imported and analysed in statistical software. A template data layout for collated data minimises the need for users to independently reformat datasets for data sharing, reducing repetition and increasing standardisation. To this end, we demonstrate the preferred NIBS-DAS data layout using the two aforementioned simulated datasets.

### ‘Long’ and ‘wide’ dataset layout

Datasets can be formatted into ‘wide’ or ‘long’ layout (Baum & Cox, 2007; Wickham, 2007; Zhang, 2016). Briefly, a ‘wide’ layout comprises one row per unique individual / entity within the dataset and separate columns per variable, whereas a ‘long’ layout contains multiple rows per individual / entity, with each row representing one measurement or timepoint (Figure 3). Datasets can be a combination of both formats, with some variables in wide format and others in long format. In the ‘Data Management’ section, we describe how dataset layouts can be manipulated using statistical software into different layouts – from wide to long (and vice versa) - depending on the analysis being performed. Example dataset layouts are demonstrated in the ‘clinical’ and ‘MEP’ datasets provided, and described below.

**Figure 3.**
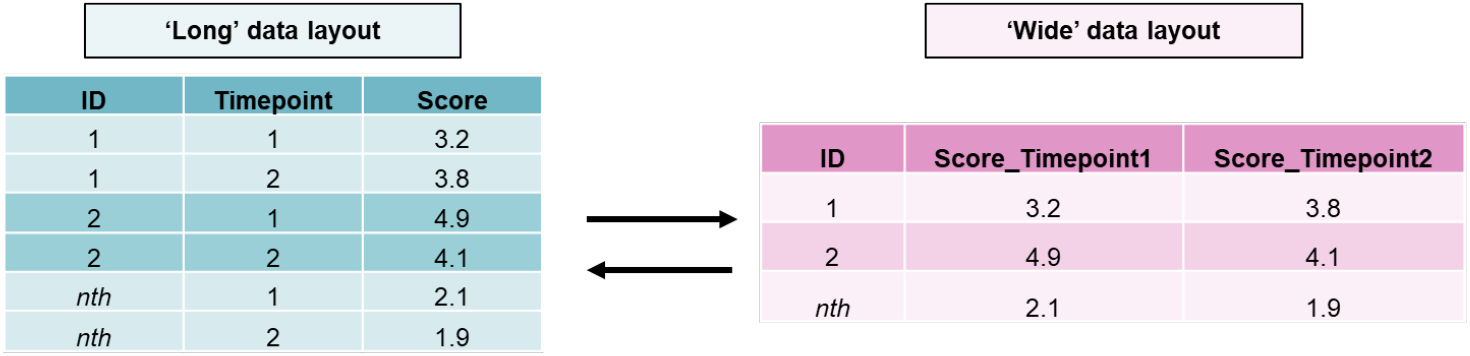
Schematic of ‘long’ (multiple rows per participant) and ‘wide’ (one row per participant) data layouts. Long formats are generally more suitable for more complex models, such as mixed-effects regression models, while wide formats are generally more suitable for more simple analyses (e.g., t-tests, one-way ANOVAs). We provide code showing how layouts can be manipulated in statistical software.

### Clinical NIBS data layout

The file ‘nibs-das_clinical_exampledata_2026-02-04.xlsx’ presents a simulated dataset comprising 60 participants, formatted such that each participant contributes two rows of data (long format) corresponding to two different measurement timepoints. The primary outcome measure is ‘Montgomery–Åsberg Depression Rating Scale’ (MADRS) symptom scores measured before and after a course of 10Hz rTMS, represented by a ‘symptom_score_madrs’ variable. Scores from other symptom scales could be added in new columns (e.g., ‘symptom_score_bdi’). The main independent variables (IVs) of this example dataset are ‘group’ (real versus sham rTMS) and ‘session_id’ (pre versus post rTMS).

### MEP data layout

The file ‘nibs-das_mep_exampledata_2026-02-04.xlsx’ shows simulated data collected using a short-interval cortical inhibition (SICI) paradigm. The dataset comprises 400 rows of data from 10 participants, with 40 MEPs from each participant (20 single-pulse MEPs, and 20 dual-pulse MEPs, evoked from the left primary motor cortex) (long format). The two MEP amplitude DVs (MEP amplitudes measured from the first dorsal interosseous [FDI] and the abductor pollicis brevis [APB] muscles) are in separate columns. In the ‘Data Management’ section, we show how to manipulate the layout of the dataset to allow for flexibility in data analysis. The main IVs are ‘tms_stim_mode’ (single versus dual MEPs) and ‘muscle’ (FDI versus APB muscles), with a ‘muscle’ variable being created when reshaping the dataset using the provided data management code files (see below).

Importantly, as much relevant data as possible should be included in collated spreadsheets, without deleting, averaging, imputing, or transforming data, which can all be done later in statistical software if necessary. For example, here all MEP amplitude observations are included in the example spreadsheet (e.g., no observations deleted, and not block averaged). This provides more flexibility in data analyses for present and future projects and increases transparency and replicability. For example, researchers may wish to look at changes in amplitudes of individual responses over time, or the factors predicting variability of MEPs (or other NIBS outcomes). For another example, if a TMS pulse fails to fire but an observation is still recorded, this observation should still be included in spreadsheets, but removed later after importing to statistical software, where it can be documented in the code files.

These practices increase transparency and prevent confusion in shared datasets.

### Data Management

This section provides a template workflow to manage data to prepare it for statistical analysis. The example clinical and neurophysiological datasets were managed using the data management (dm) code files (‘nibs-das_clinical_dm_r_2026-02-04’ and ‘nibs-das_mep_dm_r_2026-02-04’, respectively), written in R (R Core Team, 2021) (these files have also been translated to Stata (StataCorp, 2023) and are provided on our GitHub repository). Data management operations in our examples include: data importing, variable labelling and data checking, data layout manipulation ‘wide’, data layout manipulation ‘long’, appending multiple datasets, collapsing data to compute block mean values, and creating new variables. Elaboration and annotation of these operations can be found within these code files. There are many other data management decisions involved in NIBS experiments, for example, whether outlier exclusion or log transformation is performed. While these decisions are outside of the scope of the present manuscript, as above, any such operations should be documented in the code files.

Of note, there are many different types of NIBS data, and the layout of a dataset will dictate the statistical analysis that can be performed. Therefore, the NIBS-DAS pipeline shows how the layout of the original dataset can be reshaped/manipulated to provide flexibility in data analyses. For example, in the clinical example, we reshaped the ‘symptom_score_madrs’ variable into two separate columns (‘symptom_score_pre’ and ‘symptom_score_post1’) (enabling a percentage change score [‘pct_change’] in clinical symptoms), while in the MEP dataset, we show how a block mean MEP value can be computed across individual MEP observations.

Additionally, in the clinical example we demonstrate how two datasets can be combined to analyse effects across studies (i.e., ‘big data’ analyses). All data management operations and annotations can be found in the corresponding R and Stata data management files.

### Data Analysis

This section describes template/example analyses of the clinical and neurophysiological datasets. Analyses were performed using corresponding data analysis (da) code files (‘nibs-das_clinical_da_r_2026-02-04’ and ‘nibs-das_mep_da_r_2026-02-04’, respectively).

### Clinical NIBS data analysis

Here, we provide two example approaches to analyse the clinical NIBS dataset. The first analysis investigated whether there was a significant difference in pre and post rTMS symptom scores between active and sham conditions. The second analysis investigated the same question with data combined across two fictional clinical trials.

The first example (‘Example A1’ in code files) used mixed-effects linear regression with participant ID as a random intercept. Mixed-effects regression is recommended where there are multiple observations per participant, as it preserves the nesting of these non-independent datapoints (Lo & Andrews, 2015; Moen et al., 2016; Riley et al., 2010). In this mock example, there was a significant interaction between session_id and group (b = −7.90, 95% CI = −10.78, −5.02, p < 0.001) (Figure 4). Using percentage change in symptoms as the DV (Example A2) also resulted in significantly greater symptom improvement for the real versus sham rTMS group (p<0.001).

**Figure 4.**
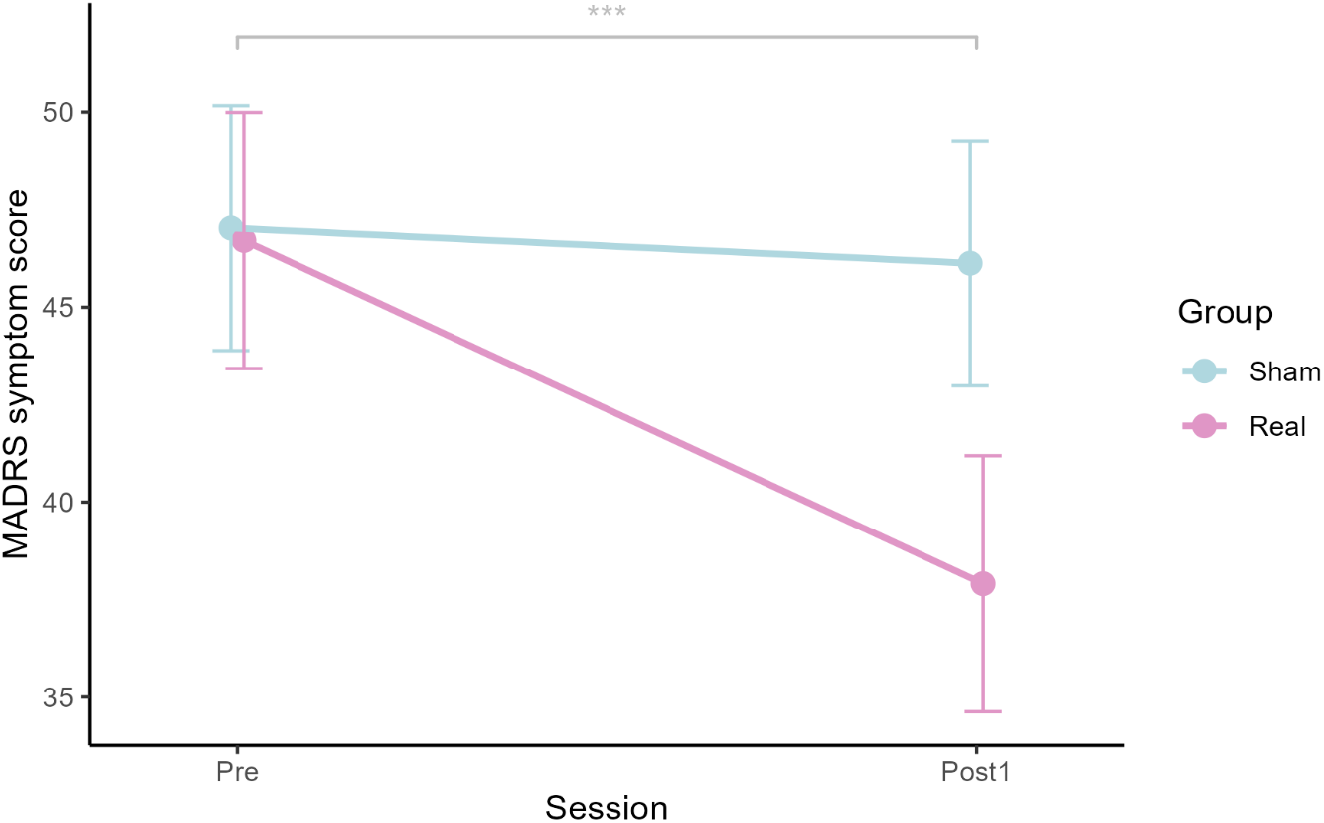
Example result from our fictional clinical rTMS experiment, demonstrating the mean values of MADRS symptom scores real (pink) and sham (blue) rTMS conditions at pre- and post-stimulation sessions.

In Example B, we demonstrate how pooled data analyses can be performed across multiple studies. Here, one can use mixed-effects regression with study ID as a random intercept (Corp et al., 2020, 2021), or as a fixed intercept where there are a low number of studies (Gomes, 2022), as in our example below. In this example, there was again a significant reduction in depression symptoms in the active condition, compared to the sham condition, across studies (b = −7.27, 95% CI = −9.87, −4.66, p < 0.001).

### MEP data analysis

In this section, we provide two example approaches to analyse the MEP dataset, both using block average MEP amplitudes as the DV. Example A investigated whether there was significant inhibition of MEPs (i.e., SICI) following left hemisphere paired-pulse TMS. This was run within the FDI and APB muscles separately, using mixed-effects regression. In both muscles, analyses demonstrated significant SICI (both p < 0.001).

After data reshaping (see Data Management section), Example B used mixed-model regression to directly compare SICI between the right FDI and APB muscles, including the created variable ‘muscle’ as an IV. Here, there was a significant interaction effect between ‘tms_stim_mode’ (single- and dual-pulse TMS) and ‘muscle’ (FDI and APB), demonstrating greater SICI in the APB, compared to the FDI, muscle (b = −0.15, 95% CI = −0.23, −0.06; p = 0.001) (Figure 5).

**Figure 5.**
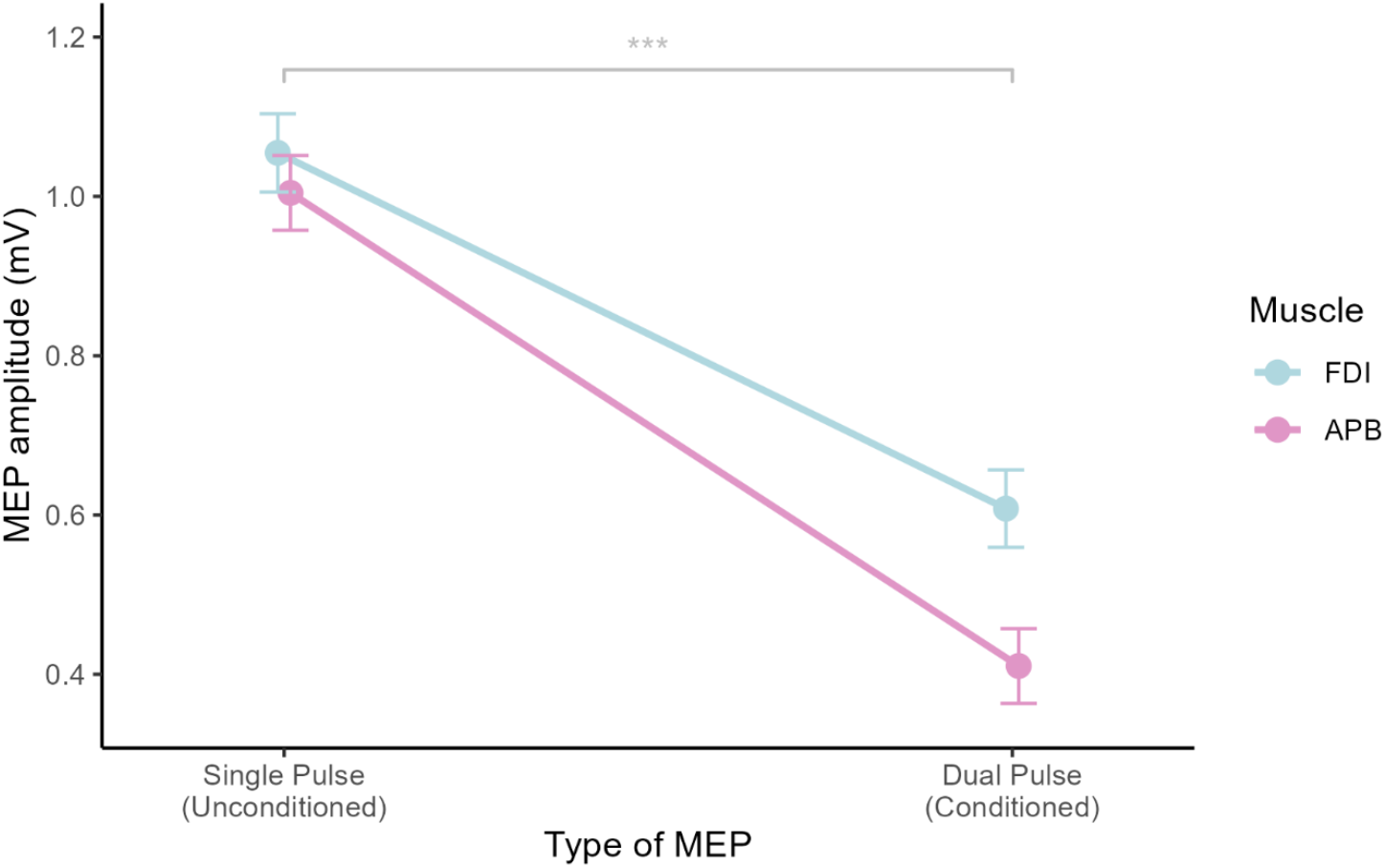
Mean MEP amplitudes for single and paired/dual pulses (i.e., unconditioned and conditioned MEPs) in the FDI (blue) and APB muscles (pink).

## Discussion

Big data approaches have been used extensively in other fields, such as MRI and neuroimaging (e.g., Karakuzu et al., 2022; Moreau et al., 2020), but their implementation in the field of NIBS has been hampered by the lack of both an accessible data sharing infrastructure and standardised guidelines for how data should be formatted for sharing. The aim of this paper was to increase the reusability and interoperability of NIBS data by creating a common data structure for how collated data should be uploaded to the Big NIBS data platform, while also providing guidelines for the data management and analysis of NIBS data. Thus, NIBS-DAS constitutes a pipeline for the analysis of collated data that can be freely used by the NIBS field in general. Since NIBS encompasses numerous techniques, such as TMS, tES, and transcranial ultrasound stimulation, NIBS-DAS is not designed as a single, regimented NIBS data structure, but rather a flexible framework that can be adapted to different NIBS techniques.

BIDS (Gorgolewski et al., 2016; Gorgolewski & Poldrack, 2016), perhaps the most common system for data management in neuroscience, is currently being extended to handle NIBS data (Bertazzoli et al., 2026), yet NIBS-BIDS will focus on the formatting of raw, machine-generated NIBS data. NIBS-DAS represents a standardised format for collated data that will expedite data sharing and reusability by reducing the need for researchers to manually re-collate raw data, which at present also leads to inconsistencies in formats. Some researchers may wish to independently re-preprocess raw data from certain NIBS methods, and extract different outcomes or use different processing methods, as is common with MRI data. However, for other types of data from NIBS studies, for example clinical trial data where the outcomes are symptom/behavioural scales, this is often unnecessary, and a collated format is highly portable and re-usable. Therefore, we anticipate that NIBS-DAS can greatly facilitate the sharing and big data analysis of NIBS clinical trial data.

A common collated data layout also facilitates the possibility for automated NIBS data pipelines in the future by providing a template schema into which raw data can be transformed. Thus, raw NIBS data could be formatted according to NIBS-BIDS guidelines and automatically collated into NIBS-DAS format. This would provide users with greater access to both raw and collated NIBS data and help form a more complete pipeline from raw NIBS data storage, through to collation, management, and analysis. We have collaborated closely with the NIBS-BIDS team on the present project to enable this possible integration in the future.

The present paper constitutes the first step towards these aims and can be considered version 1.0 of NIBS-DAS. We hope that continuing feedback will help NIBS-DAS improve in the future. Updates and future releases will be made available on *https://www.bignibsdata.com/* as they occur.

## Funding and declarations

This project has received in-kind support from Deakin University. In the last 5 years, NCR has received funding from the Australian Research Council (ARC), the Medical Research Future Fund (MRFF), the Commonwealth Scientific and Industrial Research Organisation (CSIRO), and CMAX Clinical Research PTY LTD.

## Author Contributions

**Michael P. Barham:** Conceptualisation, Methodology, Code Writing, Writing – original draft, Writing – review & editing, Project administration. **Jordan Morrison-Ham:** Methodology, Code writing, Writing – review & editing. **Christopher J. Greenwood:** Methodology, Code writing, Writing – review & editing. **Giacomo Bertazzoli:** Conceptualisation, Methodology, Code writing, Writing – review & editing. **Nigel C. Rogasch:** Conceptualisation, Methodology, Code writing, Writing – review & editing. **Hannah, G. K. Bereznicki:** Conceptualisation, Methodology, Writing – review & editing. **Ellen F. P. Younger:** Methodology, Writing – review & editing. **Elizabeth G. Ellis:** Methodology, Writing – review & editing. **Liam G. Graeme:** Methodology, Code writing, Writing – review & editing. **David A. Cunningham:** Methodology, Writing – review & editing. **Wei-Yeh Liao:** Methodology, Writing – review & editing. **Peter J. Fried:** Methodology, Writing – review & editing. **Alvaro Pascual-Leone:** Conceptualisation, Methodology, Writing – review & editing. **Peter G. Enticott:** Conceptualisation, Methodology, Writing – review & editing. **Daniel T. Corp:** Conceptualisation, Methodology, Code Writing, Writing – original draft, Writing – review & editing, Project administration.

## Acknowledgements

We would like to thank Nicholas P. Holmes for providing critical feedback on NIBS-DAS and the present manuscript.

## Notes

https://github.com/danieltcorp/nibs-das_repo

### Competing Interest Statement

The authors have declared no competing interest.

## References

Baum, C. F., & Cox, N. J. (2007). Stata Tip 45: Getting those Data into Shape. The Stata Journal, 7(2), 268–271. 10.1177/1536867X0700700211

Bertazzoli, G., Oleg, S., & Rogasch, N. (2026). NIBS-BIDS (BEP037) (Version 6) [Computer software]. https://github.com/nigelrogasch/nibsbids/tree/master/nibs-bids-v6

Bhattacharya, A., Mrudula, K., Sreepada, S. S., Sathyaprabha, T. N., Pal, P. K., Chen, R., & Udupa, K. (2022). An overview of noninvasive brain stimulation: Basic principles and clinical applications. Canadian Journal of Neurological Sciences / Journal Canadien Des Sciences Neurologiques, 49(4), 479–492. 10.1017/cjn.2021.158

Bourget, M.-H., Kamentsky, L., Ghosh, S. S., Mazzamuto, G., Lazari, A., Markiewicz, C. J., Oostenveld, R., Niso, G., Halchenko, Y. O., Lipp, I., Takerkart, S., Toussaint, P.-J., Khan, A. R., Nilsonne, G., Castelli, F. M., The BIDS Maintainers, & Cohen-Adad, J. (2022). Microscopy-BIDS: An extension to the brain imaging data structure for microscopy data. Frontiers in Neuroscience, 16, 871228. 10.3389/fnins.2022.871228

Corp, D. T., & Barham, Michael. P. (2026). Nibs-das_repo (Version 1.0) [Computer software]. https://github.com/danieltcorp/nibs-das_repo

Corp, D. T., Bereznicki, H. G. K., Barham, M. P., Clark, G. M., Chadwick, B. J., Jain, S., Khalajzadeh, H., Pascual-Leone, A., & Enticott, P. G. (2025). Big non-invasive brain stimulation data (Big NIBS data): An open-access platform and repository for NIBS data. Brain Stimulation, 18(2), 278–279. 10.1016/j.brs.2025.02.011

Corp, D. T., Bereznicki, H. G. K., Clark, G. M., Youssef, G. J., Fried, P. J., Jannati, A., Davies, C. B., Gomes-Osman, J., Kirkovski, M., Albein-Urios, N., Fitzgerald, P. B., Koch, G., Di Lazzaro, V., Pascual-Leone, A., & Enticott, P. G. (2021). Large-scale analysis of interindividual variability in single and paired-pulse TMS data. Clinical Neurophysiology, 132(10), 2639–2653. 10.1016/j.clinph.2021.06.014

Corp, D. T., Bereznicki, H. G. K., Clark, G. M., Youssef, G. J., Fried, P. J., Jannati, A., Davies, C. B., Gomes-Osman, J., Stamm, J., Chung, S. W., Bowe, S. J., Rogasch, N. C., Fitzgerald, P. B., Koch, G., Di Lazzaro, V., Pascual-Leone, A., & Enticott, P. G. (2020). Large-scale analysis of interindividual variability in theta-burst stimulation data: Results from the ‘Big TMS Data Collaboration’. Brain Stimulation, 13(5), 1476–1488. 10.1016/j.brs.2020.07.018

Gomes, D. G. E. (2022). Should I use fixed effects or random effects when I have fewer than five levels of a grouping factor in a mixed-effects model? PeerJ, 10, e12794. 10.7717/peerj.12794

Gorgolewski, K. J., Auer, T., Calhoun, V. D., Craddock, R. C., Das, S., Duff, E. P., Flandin, G., Ghosh, S. S., Glatard, T., Halchenko, Y. O., Handwerker, D. A., Hanke, M., Keator, D., Li, X., Michael, Z., Maumet, C., Nichols, B. N., Nichols, T. E., Pellman, J., … Poldrack, R. A. (2016). The brain imaging data structure, a format for organizing and describing outputs of neuroimaging experiments. Scientific Data, 3(1), 160044. 10.1038/sdata.2016.44

Gorgolewski, K. J., & Poldrack, R. A. (2016). A practical guide for improving transparency and reproducibility in neuroimaging research. PLoS Biology, 14(7), e1002506. 10.1371/journal.pbio.1002506

Guerra, A., López-Alonso, V., Cheeran, B., & Suppa, A. (2020a). Solutions for managing variability in non-invasive brain stimulation studies. Neuroscience Letters, 719, 133332. 10.1016/j.neulet.2017.12.060

Guerra, A., López-Alonso, V., Cheeran, B., & Suppa, A. (2020b). Variability in non-invasive brain stimulation studies: Reasons and results. Neuroscience Letters, 719, 133330. 10.1016/j.neulet.2017.12.058

Karakuzu, A., Appelhoff, S., Auer, T., Boudreau, M., Feingold, F., Khan, A. R., Lazari, A., Markiewicz, C., Mulder, M., Phillips, C., Salo, T., Stikov, N., Whitaker, K., & de Hollander, G. (2022). qMRI-BIDS: An extension to the brain imaging data structure for quantitative magnetic resonance imaging data. Scientific Data, 9(1), 517. 10.1038/s41597-022-01571-4

Lo, S., & Andrews, S. (2015). To transform or not to transform: Using generalized linear mixed models to analyse reaction time data. Frontiers in Psychology, 6. 10.3389/fpsyg.2015.01171

López-Alonso, V., Cheeran, B., Río-Rodríguez, D., & Fernández-del-Olmo, M. (2014). Inter-individual Variability in Response to Non-invasive Brain Stimulation Paradigms. Brain Stimulation, 7(3), 372–380. 10.1016/j.brs.2014.02.004

López-Alonso, V., Fernández-del-Olmo, M., Costantini, A., Gonzalez-Henriquez, J. J., & Cheeran, B. (2015). Intra-individual variability in the response to anodal transcranial direct current stimulation. Clinical Neurophysiology, 126(12), 2342–2347. 10.1016/j.clinph.2015.03.022

Moen, E. L., Fricano-Kugler, C. J., Luikart, B. W., & O’Malley, A. J. (2016). Analyzing clustered data: Why and how to account for multiple observations nested within a study participant? PLoS ONE, 11(1), e0146721. 10.1371/journal.pone.0146721

Moreau, C. A., Jean-Louis, M., Blair, R., Markiewicz, C. J., Turner, J. A., Calhoun, V. D., Nichols, T. E., & Pernet, C. R. (2020). The genetics-BIDS extension: Easing the search for genetic data associated with human brain imaging. GigaScience, 9(10), giaa104. 10.1093/gigascience/giaa104

Niso, G., Gorgolewski, K. J., Bock, E., Brooks, T. L., Flandin, G., Gramfort, A., Henson, R. N., Jas, M., Litvak, V., T. Moreau, J., Oostenveld, R., Schoffelen, J.-M., Tadel, F., Wexler, J., & Baillet, S. (2018). MEG-BIDS, the brain imaging data structure extended to magnetoencephalography. Scientific Data, 5(1), 180110. 10.1038/sdata.2018.110

Norgaard, M., Matheson, G. J., Hansen, H. D., Thomas, A., Searle, G., Rizzo, G., Veronese, M., Giacomel, A., Yaqub, M., Tonietto, M., Funck, T., Gillman, A., Boniface, H., Routier, A., Dalenberg, J. R., Betthauser, T., Feingold, F., Markiewicz, C. J., Gorgolewski, K. J., … Ganz, M. (2022). PET-BIDS, an extension to the brain imaging data structure for positron emission tomography. Scientific Data, 9(1), 65. 10.1038/s41597-022-01164-1

Pernet, C. R., Appelhoff, S., Gorgolewski, K. J., Flandin, G., Phillips, C., Delorme, A., & Oostenveld, R. (2019). EEG-BIDS, an extension to the brain imaging data structure for electroencephalography. Scientific Data, 6(1), 103. 10.1038/s41597-019-0104-8

Polanía, R., Nitsche, M. A., & Ruff, C. C. (2018). Studying and modifying brain function with non-invasive brain stimulation. Nature Neuroscience, 21(2), 174–187. 10.1038/s41593-017-0054-4

R Core Team. (2021). R: A language and environment for statistical computing [Computer software]. R Foundation for Statistical Computing. https://www.R-project.org/

Riley, R. D., Lambert, P. C., & Abo-Zaid, G. (2010). Meta-analysis of individual participant data: Rationale, conduct, and reporting. BMJ, 340, c221. 10.1136/bmj.c221

StataCorp, L. (2023). Stata statistical software: Release 18 [Computer software]. College Station, TX. https://www.stata.com/stata18/

Wickham, H. (2007). Reshaping Data with the reshape Package. Journal of Statistical Software, 21, 1–20. 10.18637/jss.v021.i12

Wilkinson, M. D., Dumontier, M., Aalbersberg, Ij. J., Appleton, G., Axton, M., Baak, A., Blomberg, N., Boiten, J.-W., Da Silva Santos, L. B., Bourne, P. E., Bouwman, J., Brookes, A. J., Clark, T., Crosas, M., Dillo, I., Dumon, O., Edmunds, S., Evelo, C. T., Finkers, R., … Mons, B. (2016). The FAIR Guiding Principles for scientific data management and stewardship. Scientific Data, 3(1), 160018. 10.1038/sdata.2016.18

Zhang, Z. (2016). Reshaping and aggregating data: An introduction to reshape package. Annals of Translational Medicine, 4(4), 78. 10.3978/j.issn.2305-5839.2016.01.33

